# Regionally Diffuse Muscle Pain-Hypersensitivity in Humans During Acute Muscle Pain

**DOI:** 10.1101/525832

**Authors:** J. S. Dunn, S. S. Nagi, D. A. Mahns

## Abstract

**Background:** We have previously shown that an intramuscular infusion of 5% hypertonic saline (HS) produces a painful response to normally innocuous stimuli applied to overlying and adjacent *skin* regions. In the current study, we explored whether a similar interaction could be observed between adjacent, contralateral and remote muscles. Indeed, widespread muscle pain-hypersensitivity is a hallmark of chronic pain conditions such as fibromyalgia.

**Methods:** 5% HS was infused into the *flexor carpi ulnaris* (FCU) muscle to develop a stable baseline pain (n=30). In separate experiments, each of the three test locations (n=10 per site), the adjacent *abductor digiti minimi* (ADM), contralateral FCU and contralateral *tibialis anterior* (TA) (**part 1-3**, respectively), 50μL of 0.9% normal saline (NS) was infused (in triplicate) prior to, during and following HS-induced muscle pain.

**Results:** Under control conditions (no background pain), the infusion of NS was *imperceptible* by all subjects. In the presence of HS-induced background pain (FCU), in ***part 1*** the NS co-infusion into ADM increased overall pain by 17%. This was replicated in the contralateral FCU (***part 2***) with a 12% pain increase, and in the TA (***part 3***) with a 15% pain increase in response to the NS co-infusions. Notably, over 80% of subjects perceived the NS-induced increase in pain at the HS-infusion location (FCU) rather than the NS-infusion location (adjacent, contralateral and remote).

**Conclusions:** Intramuscular infusion of HS results in pain-hypersensitivity to sub-perceptual stimulation of muscle afferents in a somatotopically unrestricted manner, indicating the involvement of a central (likely supra-spinal) mechanism.

**Significance:** This work provides evidence for a regionally diffuse type of pain hypersensitivity, manifesting as a painful response to normally sub-perceptual stimulation in the context of acute experimentally induced muscle pain. This phenomenon may provide parallels to clinically relevant painful conditions and neuropathies.

## Introduction

For most individuals it is relatively easy to distinguish between innocuous and noxious stimuli. However, in a subset of individuals afflicted by chronic pain there is a disturbance of normal somatosensory function, such that a normally innocuous stimulus can evoke pain (Berglund et al., 2002; Clauw, 2014; Wolfe et al., 1995). Such pain manifestations have debilitating impacts on both the individual (Clauw, 2014; Koroschetz et al., 2011) and society (Doth et al., 2010; van Leeuwen et al., 2012).

It has been observed that inputs from large-diameter (Aβ/Group I-II) mechano-sensitive afferents in the skin and muscle can contribute to pain hypersensitivity in acute and chronic pain states (Price et al., 1992; Torebjörk et al., 1992; Weerakkody et al., 2003; Weerakkody et al., 2001). It is postulated that this phenomenon involves the convergence of inputs from superficially terminating nociceptive small-diameter fibres and the deeper terminating mechanoreceptive large-diameter fibres in the dorsal horn (Basbaum et al., 2009; Brown, 1982; Purves et al., 2001). In addition to the role of large-diameter mechano-afferents in pain hypersensitivity, an increasing body of evidence has implicated their unmyelinated counterparts, C-tactile fibres in the skin (Nagi et al., 2015; Nagi et al., 2011; Samour et al., 2015; Seal et al., 2009). Furthermore, such studies have revealed that hypertonic saline (HS)-evoked pain induces a state of touch-evoked pain (allodynia) that extends to overlying (Nagi et al., 2011; Samour et al., 2015) and adjacent (distal) skin regions (Nagi & Mahns, 2013; Nagi et al., 2011; Samour et al., 2015). Likewise, intramuscular HS produces a deep musculoskeletal pain that often extends or refers to distal regions (Graven-Nielsen, 2006; Graven-Nielsen et al., 1997a, 1997b; Kellgren, 1938; Steinbrocker et al., 1953), produces a remapping of cortical pain areas (Rubin et al., 2010) and a pain hypersensitivity that extends bilaterally (Samour et al., 2017). These complex interactions cannot readily be explained by changes in peripheral circuitry and appear to mimic characteristics of chronic pain conditions such as fibromyalgia. Within such chronic conditions, current arguments favour an explanation based on a central change in, or sensitization of, neural function that results in the observed widespread and diffuse musculoskeletal pain, pressure-pain hypersensitivity, cutaneous allodynia and tactile dysesthesia (Ablin et al., 2008; Berglund et al., 2002; Bradley, 2009; Case et al., 2016; Clauw, 2009).

In this study a HS infusion model was used in order to examine whether the interaction previously observed between muscle and skin (Nagi & Mahns, 2013; Nagi et al., 2011; Samour et al., 2015; Samour et al., 2017) could be replicated between adjacent and remote muscles. We hypothesized that the presence of background (sustained) nociceptive activity using HS infusion would result in a state of central sensitization, such that the application of a normally innocuous stimulus (normal saline, NS) subsequently would result in exacerbation of overall pain, regardless of whether the NS was infused into adjacent or remote muscles.

## Methods

24 healthy subjects aged 18-28 years (6 females), with no reported history of musculoskeletal or neurological disorders, were recruited for this study. Subjects were also asked to abstain from intensive bouts of exercise for 48 hours preceding the experiment so as not to sensitize the target muscles (Weerakkody et al., 2001). Eight subjects participated in multiple arms of the study across different experimental sittings. Informed written consent was obtained from each participant prior to each experiment. This study was approved by the Human Research Ethics Committee (approval number: H9190) of Western Sydney University in accordance with the revised Declaration of Helsinki.

Subjects sat comfortably in a chair throughout all experiments. Infusions of HS and NS were performed using a Harvard Apparatus Syringe Infusion Pump 22 (Harvard Apparatus, South Natick, Massachusetts, USA). Pain ratings were continuously recorded using an ADInstruments Response meter with input run through an ADInstruments Power Lab (ADInstruments, Dunedin, New Zealand).

### Infusion of hypertonic saline

Across all arms of the study, 5% HS was infused into the belly of the *flexor carpi ulnaris* (FCU) muscle for ~10 min to ensure for the baseline pain to stabilize. The infusion rate of HS in the FCU varied between subjects (30-175 μL/min) in order to establish a moderate pain intensity between 4 and 6 on a visual analog scale (VAS) ranging from 0 (no pain) to 10 (worst pain).

### Infusion of normal saline

After a stable baseline pain was maintained for at least a minute, HS-NS co-infusion events followed. NS (0.9%) was infused at the rate of 50 μL/min for 1 min per trial (tested in triplicate). This duration was chosen based on the data collected in a pilot study which indicated a delay of ~30 s before the onset of an increase in pain levels. Subjects were asked to rate the overall pain intensity, and any changes thereof, on the VAS. Care was taken to avoid the use of suggestive language before the subjects. The triplicate NS trials were performed at ~1-min intervals. During these intervals, subjects were asked to verbally localize the region of pain localisation.

In addition to concurrent HS-S infusions, NS alone was infused in triplicate trials prior to the commencement and upon cessation of HS-evoked pain in all experiments. Typically, the HS-evoked pain disappeared over a time course of under 10 min. After a 3-5 min wait following cessation of pain, NS infusion was repeated at each site. Collectively, ≈450 μL of NS was infused per muscle.

### Part 1: Interactions with adjacent muscles

NS was infused into the ipsilateral *abductor digiti minimi* (ADM) in order to examine potential interactions between *adjacent* muscles in response to HS induced acute muscular pain. The ADM muscle was chosen as it shares the same peripheral innervation (ulnar nerve) as the HS infused FCU.

### Part 2: Contralateral interactions

NS was infused into the belly of contralateral FCU muscle in order to test whether the HS-NS interactions were limited to muscles within the same nerve territory or spread to contralateral muscles as well (i.e. central involvement).

### Part 3: Remote interactions

NS was delivered to the belly of the *tibialis anterior* (TA) muscle in order to determine the spatial extent of inter-muscle interactions in an acute pain state.

### Statistical analysis

Repeated measures two-way analysis of variance (RM 2-way ANOVA) was used to compare pain evoked at baseline (HS infusion alone) with test responses (co-infusion of NS and HS) at each location (adjacent, contralateral and remote). Where a significant change (p < 0.05) was found, individual comparisons were made using a Tukey’s multiple comparison test. The normal distribution of data was confirmed in all groups using D’Agostino and Pearson omnibus normality test. Pain scores for the baseline (HS) and co-infusion (HS and NS) conditions are presented as mean ± standard error of the mean (SEM) for all parts of the study. Statistical analysis was performed using GraphPad Prism 7.04 software (La Jolla, California, USA).

## Results

Prior to the induction and following the cessation of HS evoked muscle pain, all subjects reported NS infusion (50 μL/min) to be imperceptible (i.e. VAS=0 with no associated percept). In contrast, during the infusion of 5% HS into the FCU always resulted in a diffuse, deep pain in the muscle that extended down the medial aspect of the forearm and remained stable in the absence of concurrent NS-infusions (see Figure 1A as example). At all three test locations (adjacent, contralateral and remote), infusion of NS significantly increased the HS-evoked pain in all trials (T1-3, p<0.05, Figure 1B-D left hand panel). The pooled (*n*=3) mean response for all 10 subjects, with respective HS and HS+NS data points linked, are shown in the right hand panel of Figure 1B-D (p<0.0001). At each test location, pain returned to baseline (HS) within 1-min of cessation of NS-co-infusion. The stability of Baseline and reproducibility of co-infusion responses were confirmed by individual RM 2-way ANOVA for each location with individual differences confirmed by Tukey’s multiple comparisons test; (see below)

**Figure 1.**
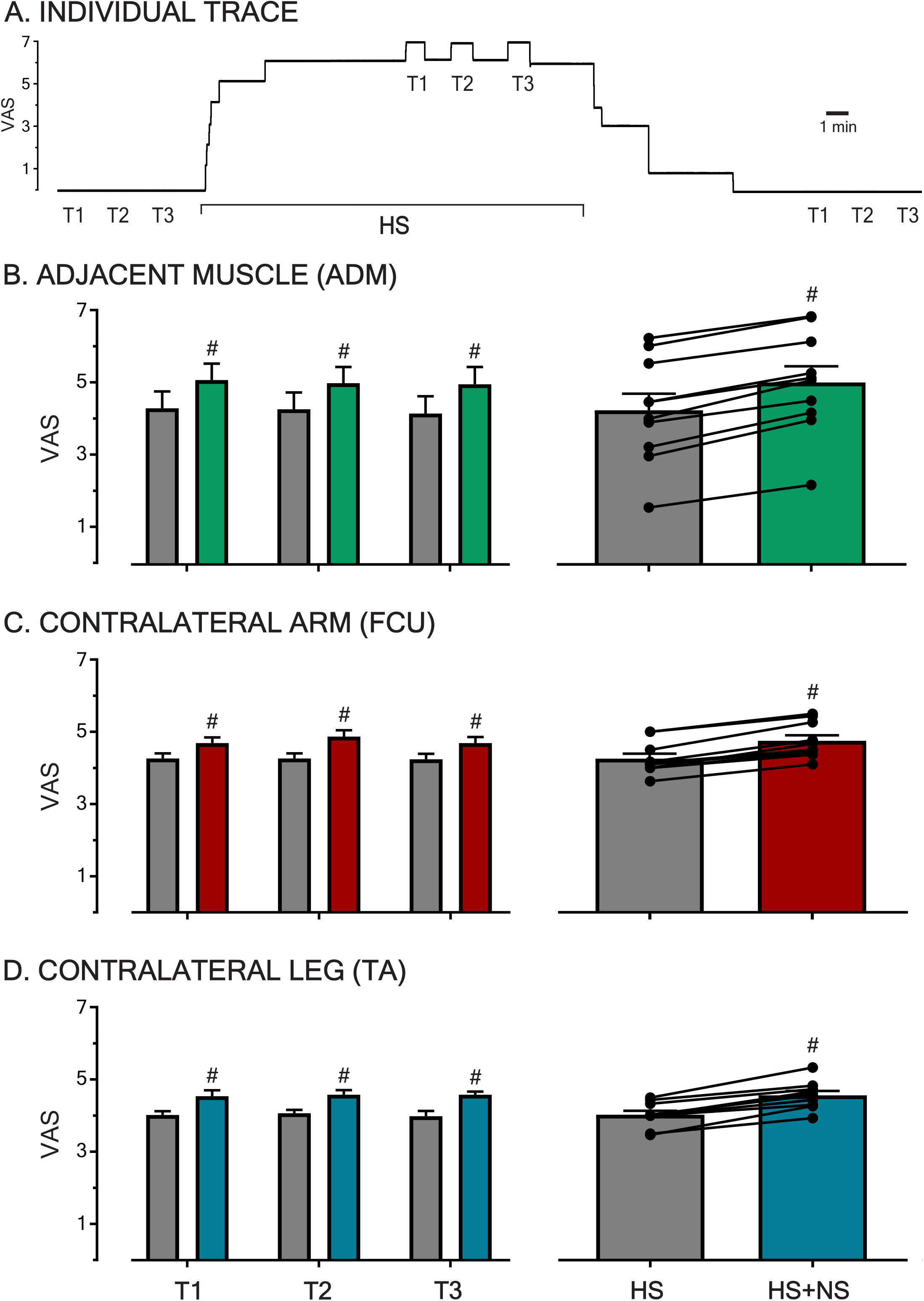
Pain intensities as reported on VAS in response to HS infusion and subsequent to transient NS co-infusion trials at various sites across the body. **A.** A raw trace of a subject’s VAS ratings throughout an experimental sitting is shown. In the absence of background pain infusions of NS for 1 min (T1, T2, T3) were imperceptible. During baseline HS induced muscle pain in the FCU, co-infusion of NS (triplicate, left panel B-D) produced a reproducible increase in muscle pain. Following cessation of HS infusion and the associated background pain (VAS=0) NS trials were once again imperceptible. In all three sessions HS pain was generated in the FCU, the test location for NS infusion were the *adjacent* ADM muscle (**B**) and the *contralateral* FCU (**C**) and TA muscles (**D**, *remote*). At each test location, co-infusion with the previously imperceptible NS during HS background pain resulted in a reproducible (n=3) and significant increase in muscle pain (p<0.0001, right panel B-D). The transient pain increase was reproducible across trials at all sites. Significant changes (p<0.0001, #) were confirmed between baseline (HS) and co-infusion (HS+NS) using RM 2-way ANOVA.

### Part 1: Interactions in adjacent muscles

In this arm of the study, the infusion of HS into the FCU resulted in an average pain score of 4.3±0.5 on the VAS (*n*=10). When NS was co-infused into the adjacent ADM in the presence of this background pain, the overall pain score increased to 5.0±0.4 (Figure 1B). This constitutes an average increase in pain of 17% and when comparing the baseline and co-infusion pain scores this pain increase is found to be significant using a RM 2-way ANOVA (p<0.0001, F (1,27) = 318.5). Furthermore, whilst the infusion of NS in the ADM caused this significant increase in overall pain, this pain increase was localised to the FCU in eight out of ten subjects, with no discernible percept attributed to the ADM. This indicates that overall muscle pain can be modulated in a reproducible and stimulus-locked manner by repeated sub-perceptual stimulation of an adjacent muscle.

### Part 2: Contralateral interactions

The infusion of HS in the FCU (*n*=10) resulted in an average VAS score of 4.3±0.1. The co-infusion of NS in the contralateral FCU increased the overall pain score to 4.8 ±0.2 (Figure 1C). This represents a 12% increase in subject pain scores during co-infusion, an effect found to be significant (p<0.0001, F (1,27) = 156.7). Once again, despite the infusion of NS into the contralateral limb, an increase in pain was perceived at the site of HS infusion in the FCU whilst the NS infusion remained imperceptible in nine out of ten subjects. The demonstration that these interactions are not limited to adjacent regions, but can be elicited across contralateral muscles, suggests a central underpinning to this phenomenon.

### Part 3: Remote interactions

Within this aspect of the study, subjects reported an average pain level of 4.0 ±0.1 (n=10) to infusion of HS in the FCU. When NS was administered to the contralateral TA, a pain increase of 15% was reported with the overall pain intensity increasing to 4.6±0.1 (Figure 1D). A comparison of baseline and co-infusion VAS scores revealed a significant increase (p<0.0001, F (1,27) = 97.84). This pain increase was not felt at the site of NS infusion in the TA, but rather as an increase in pain at the site of HS infusion in the FCU of the contralateral arm in nine out of ten subjects. The observed interaction between the site of noxious muscle stimulation and remote innocuous muscle stimulation alludes to the involvement of a supra-spinal mechanism.

In Figure 2, triplicate responses for each individual (n=10) at all three test sites (n=90), to transient NS infusion during HS infusion (i.e. HS+NS) have been plotted as a function of the baseline pain evoked by HS alone. When plotted in this manner all data points fell to the left of the line of equivalence (45° line) indicating that the NS infusion evoked a reproducible pain increase across the entirety (VAS 1.4-6.7) of baseline pain tests, a range that was defined by the FCU-ADM group.

**Figure 2.**
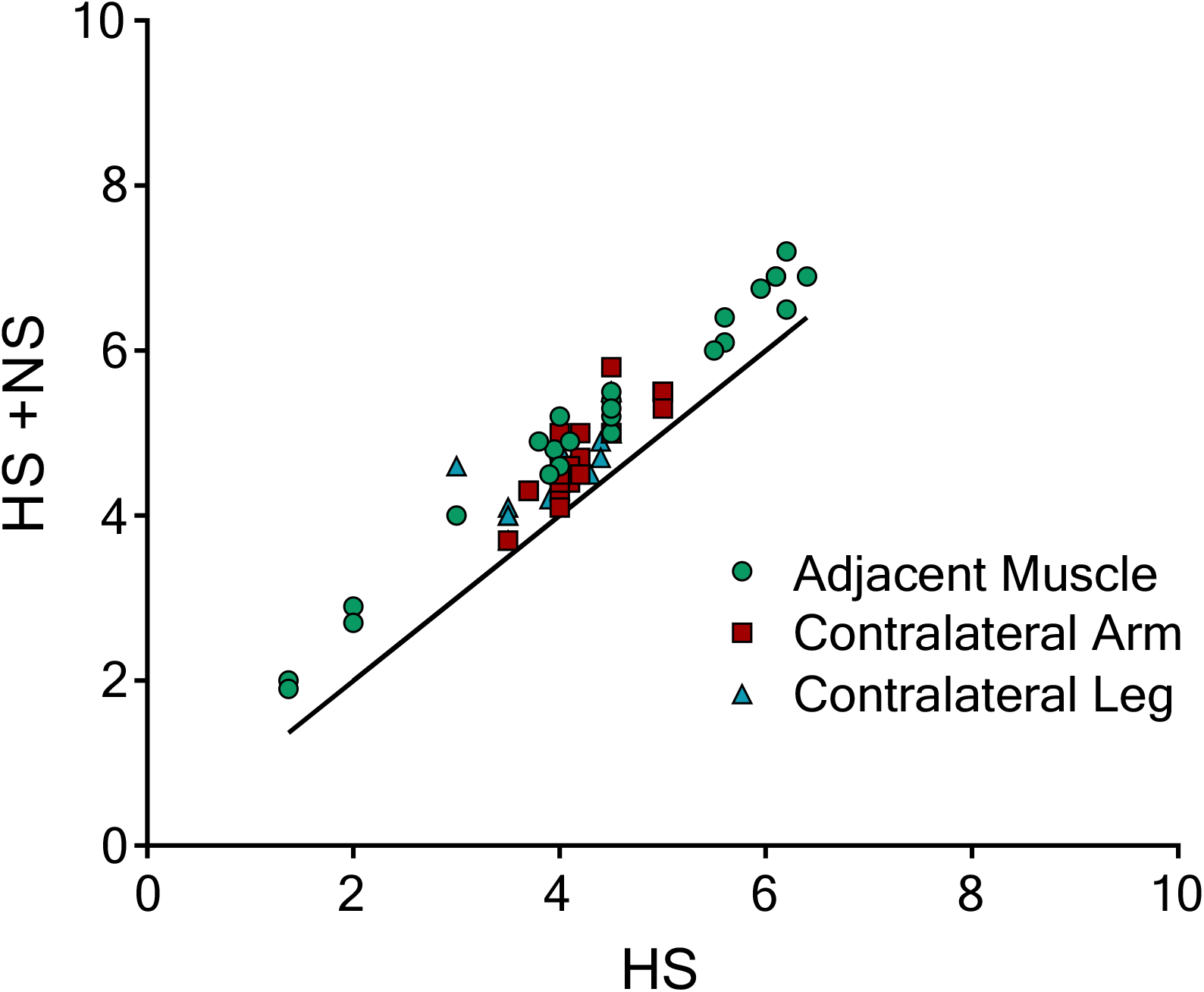
Triplicate data points for each subject during HS-NS co-infusion plotted as a function of baseline pain. When triplicate responses from each individual (n=10) at each location (n=90) to NS infusion during HS infusion (HS+NS) are plotted as a function of baseline pain (i.e. HS alone) all data points fall to the right of the line of equivalence. This indicates that the NS infusion evoked a reproducible effect between trials and across all test sites.

## Discussion

The current study has provided evidence that during HS-induced muscle pain in the arm (FCU), muscular hypersensitivity extents to adjacent muscles (ADM), the contralateral arm (FCU), and contralateral leg (TA) in a somatotopically unrestricted manner. This finding not only builds upon the previous observation that an intramuscular HS infusion can result in allodynia in the overlying and adjacent skin regions (Nagi & Mahns, 2013; Nagi et al., 2011; Samour et al., 2015) but reinforces the role of the central nervous system (CNS) in underpinning this phenomenon.

The sub-perceptual nature of repeated intermittent infusions (50 μL over 1 min) under control conditions suggests that localised muscle distension need not activate the nociceptors, and may well activate low-threshold stretch-sensitive receptors within muscle. In this respect, these weak mechanical stimuli resemble the inability of weak (micro) intraneural electrical stimulation to produce a discernible pain sensation at recording sites dominated by muscle spindles (Gandevia, 1985; Macefield et al., 1990). The conversion of the sub-perceptual NS stimulus to one that enhances pain, during HS infusion in the FCU muscle, is unlikely to be due to peripheral sensitization given the anatomical separation (arm vs hand, >15 cm) and small volume of intermittently infused NS. Likewise, the corresponding pain evoked from the contralateral arm is more consistent with a central involvement. Furthermore, the interaction between the FCU and the contralateral TA suggests that the central involvement likely extends to supra-spinal structures. This broad ranging interaction appears to be in marked contrast to the somatotopically constrained interactions observed in the skin; for example, the confinement of secondary hyperalgesia to the region immediately surrounding intradermal capsaicin injection (Ali et al., 1996; Magerl et al., 1998; Simone et al., 1989) or the inability of microstimulation of large-diameter mechanoreceptors innervating a skin region beyond the site of pain hyperalgesia to produce a painful percept (Torebjörk et al., 1992).

The use of sub-perceptual stimuli during muscle pain revealed a profoundly limited differentiation of pain locognosia, such that >80% of subjects reported that pain increased at the HS site (FCU) when NS was infused into adjacent (ADM), contralateral (FCU) and remote (TA) sites. The NS induced pain increase was time-locked to the transient NS infusion and was referred to the HS site despite the subject being well aware of needle insertion (prior to test commencement) in the adjacent, contralateral or remote muscle. Assertions as to the exact location of this CNS involvement in mediating this diffuse hypersensitivity cannot be resolved by this study. Nonetheless, the observation of a regionally diffuse pain-hypersensitivity in an acute pain model demonstrates that the requisite central circuitry may already be present, and thus an elaborate anatomical reorganisation need not be necessary for this to occur.

The diffuse pain-hypersensitivity observed in the current study is most likely driven by a transient and reversible episode of central sensitization (Samour et al., 2017) in response to the HS-induced muscle pain. The HS infusion alone was run for ~10 min prior to the commencement of NS co-infusion. The nociceptive input during this period may have allowed for sensitization of the wide-dynamic-range (WDR) neurons in the dorsal horn (Arendt-Nielsen & Henriksson, 2007; Yunus, 2007) and thus a state of central sensitization to develop. Intriguingly, recent molecular work in animal models has shown that peripheral nociceptive signalling can lead to interactions in pain processing pathways right up to the cortex (Tochiki et al., 2016). Indeed, the clinical correlates of central sensitization (Ablin et al., 2008; Yunus, 2007) are apparent in a HS infusion model with hyperalgesia and allodynia reported in this and previous work (Nagi & Mahns, 2013; Nagi et al., 2011; Samour et al., 2015; Samour et al., 2017).

In regards to the experimental model of intramuscular HS infusion itself, this study is novel in its findings that an acute intramuscular HS infusion can result in adjacent (ipsilateral), contralateral and remote muscular pain-hypersensitivities to otherwise imperceptive stimuli. Previous findings have shown that repeated intramuscular HS injections in the TA results in a pressure-pain hypersensitivity developing across both the ipsilateral and contralateral TA muscles (Samour et al., 2017). The convergent evidence suggests the formation of regionally diffuse sensitivities in HS-induced pain models. It should now be considered that a contralateral or even remote limb should not be used as a ‘control’ sample as somatotopically unrestrained hypersensitivity may have manifested as a result of the painful intervention (Shaikh et al., 2016), confounding any data obtained. Rather, the evidence for centralized effects of intramuscular HS necessitates the need for control data collection prior to any HS administration, and warrants investigation in other commonly used pain models.

The demonstration of certain clinical correlates of chronic pain conditions in the HS-infusion model, such as widespread muscular pain hypersensitivity and cutaneous allodynia which is often reported in fibromyalgia (Berglund et al., 2002; Bradley, 2009; Clauw, 2009; Gracely et al., 2003), may provide an experimental model for these conditions with the distinct advantage of being transient in nature with full return to normalcy upon cessation of infusion.

## Conclusions

Overall, it is evident from the data that the infusion of HS into a muscle results in a centralized hypersensitivity which evokes an exacerbation of HS-induced pain from sub-perceptual stimulation of adjacent and remote muscles in a somatotopically unconstrained manner. Further study of the mechanisms behind the diffuse hypersensitivity seen in the HS infusion model may advance our understanding of the conditions characterised by chronic musculoskeletal pain including fibromyalgia.

## Acknowledgments

The authors would like to acknowledge the work of Sophia Mavromihalis in the collection of pilot data for this project.

## Author contributions

The original study was conceived by S. S. Nagi and D. A. Mahns. J. S. Dunn and S. S. Nagi carried out all experiments. All authors contributed to the design of the study and discussed the results before the first drafting of the manuscript by J. S Dunn. D. A. Mahns and S. S. Nagi contributed to subsequent revisions. All authors read and approved the final manuscript prior to submission.

## Notes

<u>Funding Sources</u>: This work was supported by the Western Sydney University School of Medicine. The funding body had *no* role in the design of the study and collection, analysis, and interpretation of data and in manuscript preparation.

<u>Disclosures</u>: The authors declare no conflicts of interest.

## References

Ablin, J., Neumann, L., & Buskila, D. (2008). Pathogenesis of fibromyalgia - A review. Joint Bone Spine, 75, 273–279.

Ali, Z., Meyer, R. A., & Campbell, J. N. (1996). Secondary hyperalgesia to mechanical but not heat stimuli following a capsaicin injection in hairy skin. Pain, 68, 401–411.

Arendt-Nielsen, L., & Henriksson, K. G. (2007). Pathophysiological mechanisms in chronic musculoskeletal pain (fibromyalgia): the role of central and peripheral sensitization and pain disinhibition. Best Pract Res Clin Rheumatol., 21(3), 465–480.

Basbaum, A. I., Bautista, D. M., Scherrer, G., & Julius, D. (2009). Cellular and molecular mechanisms of pain. Cell, 139, 267–289.

Berglund, B., Harju, E., Kosek, E., & Lindblom, U. (2002). Quantitative and qualitative perceptual analysis of cold dysesthesia and hyperalgesia in fibromyalgia. Pain, 96, 177–187.

Bradley, L. A. (2009). Pathophysiology of Fibromyalgia. Am. J. Med, 122, S22–S30.

Brown, A. G. (1982). The dorsal horn of the spinal cord. Q. J. Exp. Physiol., 67, 193–212.

Case, L. K., Čeko, M., Gracely, J. L., Richards, E. A., Olausson, H., & Bushnell, M. C. (2016). Touch perception altered by chronic pain and by opiod blockade. eNeuro, 10(3). Retrieved from http://www.eneuro.org/content/eneuro/3/1/ENEUR0.0138-15.2016.full.pdf doi:10.1523/ENEURO.0138-15.2016

Clauw, D. J. (2009). Fibromyalgia: An overview. Am. J. Med, 122, S3–S13.

Clauw, D. J. (2014). Fibromyalgia - A clinical review. JAMA, 311(15), 1547–1555.

Doth, A. H., Hansson, P. T., Jensen, M. P., & Taylor, R. S. (2010). The burden of neuropathic pain: a systematic review and meta-analysis of health utilities. Pain, 149(2), 338–344. doi:10.1016/j.pain.2010.02.034

Gandevia, S. C. (1985). Illusory movements produced by electrical stimulation of low-threshold muscle afferents from the hand. Brain, 108 (Pt 4), 965–981.

Gracely, R. H., Grant, M. A. B., & Giesecke, T. (2003). Evoked pain measures in fibromyalgia. Best Pract Res Clin Rheumatol., 17(4), 593–609.

Graven-Nielsen, T. (2006). Fundamentals of muscle pain, referred pain, and deep tissue hyperalgesia. Scand J Rheumatol Suppl, 35(122), 1–43.

Graven-Nielsen, T., Arendt-Nielsen, L., Svensson, P., & Jensen, T. S. (1997a). Experimental muscle pain: A quantitative study of local and referred pain in humans following injection of hypertonic saline. J Musculoskelet Pain, 5(1), 49–69.

Graven-Nielsen, T., Arendt-Nielsen, L., Svensson, P., & Jensen, T. S. (1997b). Quantification of local and referred muscle pain in humans after sequential i.m. injections of hypertonic saline. Pain, 69(1-2), 111–117.

Kellgren, J. H. (1938). Observations on referred pain arising from muscle. Clin Sci, 3, 175–190.

Koroschetz, J., Rehm, S. E., Gockel, U., Brosz, M., Freynhagen, R., Tölle, T. R., & Baron, R. (2011). Fibromyalgia and neuropathic pain - differences and similarities. A comparison of 3057 patients with diabetic painful neuropathy and fibromyalgia. BMC Neurol, 11(1), 55.

Macefield, G., Gandevia, S. C., & Burke, D. (1990). Perceptual responses to microstimulation of single afferents innervating joints, muscles and skin of the human hand. J. Physiol., 429(1), 113–129.

Magerl, W., Wilk, S. H., & Treede, R. D. (1998). Secondary hyperalgesia and perceptual wind-up following intradermal injection of capsaicin in humans. Pain, 74, 257–268.

Nagi, S. S., Dunn, J. S., Birznieks, I., Vickery, R. M., & Mahns, D. A. (2015). The effects of preferential A- and C-fibre blocks and T-type calcium channel antagonist on detection of low-force monofilaments in healthy human participants. BMC Neuroscience, 16(1), 1–10. doi:10.1186/s12868-015-0190-2

Nagi, S. S., & Mahns, D. A. (2013). Mechanical allodynia in human glabrous skin mediated by low-threshold cutaneous mechanoreceptors with unmyelinated fibres. Exp Brain Res, 231, 139–151.

Nagi, S. S., Rubin, T. K., Chelvanayagam, D. K., Macefield, V. G., & Mahns, D. A. (2011). Allodynia mediated by C-tactile afferents in human hairy skin. J. Physiol., 589, 4065–4075.

Price, D., Long, S., & Huitt, C. (1992). Sensory testing of pathophysiological mechanisms of pain in patients with reflex sympathetic dystrophy. Pain, 49, 163–173.

Purves, D., Augustine, G. J., Fitzpatrick, D., Katz, L. C., LaMantia, A., McNamara, J. O., & Williams, S. M. (2001). The Major Afferent Pathway for Mechanosensory Information: The Dorsal Column-Medial Lemniscus System. In Neuroscience (2 ed.). Sunderland: Sinauer Associates.

Rubin, T. K., Henderson, L. A., & Macefield, V. G. (2010). Changes in the spatiotemporal expression of local and referred pain following repeated intramuscular injections of hypertonic saline: a longitudinal study. J Pain, 11(8), 737–745.

Samour, M. S., Nagi, S. S., & Mahns, D. A. (2015). Cav3.2-expressing low-threshold C fibres in human hairy skin contribute to cold allodynia - a non-TRPV1 and non-TRPM8-dependent phenomenon. Pain, in press.

Samour, M. S., Nagi, S. S., Shortland, P. J., & Mahns, D. A. (2017). Minocycline prevents muscular pain hypersensitivity and cutaneous allodynia produced by repeated intramusuclar injections of hypertonic saline in healthy human participants. J Pain, 18(8), 994–1005.

Seal, R. P., Wang, X., Guan, Y., Raja, S. N., Woodbury, J., Basbaum, A. I., & Edwards, R. H. (2009). Injury-induced mechanical hypersensitivity requires C-low threshold mechanoreceptors. Nature, 462, 651–655.

Shaikh, S., Shortland, P., Lauto, A., Barton, M., Morley, J. W., & Mahns, D. A. (2016). Sensory perturbations using suture and sutureless repair of transected median nerve in rats. Somatosens Mot Res, 33(1), 20–28. doi:10.3109/08990220.2016.1142438

Simone, D. A., Baumann, T. K., & LaMotte, R. H. (1989). Dose-dependent pain and mechanical hyperalgesia in humans after intradermal injection of capsaicin. Pain, 38, 99–107.

Steinbrocker, O., Isenberg, S. A., Silver, M., Neustadt, D., Kuhn, P., & Schittone, M. (1953). Observations on pain produced by injection of hypertonic saline into muscles and other supportive tissues. J Clin Invest., 32(10), 1045–1051.

Tochiki, K. K., Maiarú, M., Norris, C., Hunt, S. P., & Géranton, S. M. (2016). The mitogen and stress-activated protein kinase 1 regulates the rapid epigenetic tagging of dorsal horn neurons and nocifensive behaviour. Pain, 157, 2594–2604.

Torebjörk, H. E., Lundberg, L. E. R., & LaMotte, C. (1992). Central changes in processing of mechanoreceptive input in capsaicin-induced secondary hyperalgesia in humans. J. Physiol., 448, 765–780.

van Leeuwen, M. T., Blyth, F. M., March, L. M., Nicholas, M. K., & Cousins, M. J. (2012). Chronic pain and reduced work effectiveness: The hidden cost to Australian employers. Eur. J. Pain, 10(2), 161–166.

Weerakkody, N. S., Percival, P., Hickey, M. W., Morgan, D. L., Gregory, J. E., Canny, B. J., & Proske, U. (2003). Effects of local pressure and vibration on muscle pain from eccentric exercise and hypertonic saline. Pain, 105, 425–435.

Weerakkody, N. S., Whitehead, N. P., Canny, B. J., Gregory, J. E., & Proske, U. (2001). Large-fiber mechanoreceptors contribute to muscle soreness after eccentric exercise. J Pain, 2(4), 209–219.

Wolfe, F., Ross, K., Anderson, J. M., Russell, I. J., & Hebert, L. (1995). The prevalence and characteristics of fibromyalgia in the general population. Arthritis Rheum, 38(1), 19–28.

Yunus, M. B. (2007). Role of central senitization in symptoms beyond muscle pain, and the evaluation of a patient with widespread pain. Best Pract Res Clin Rheumatol., 21(3), 481–497.

